# Decoupling diet from microbiome dynamics results in model mis-specification that implicitly annuls potential associations between the microbiome and disease phenotypes—ruling out any role of the microbiome in autism (Yap *et al.* 2021) likely a premature conclusion

**DOI:** 10.1101/2022.02.25.482051

**Authors:** James T. Morton, Sharon M. Donovan, Gaspar Taroncher-Oldenburg

**Affiliations:** Simons Foundation; University of Illinois

## Abstract

Recently, and in a tour de force effort, Yap *et al*. performed a comprehensive association analysis of factors such as demographics, psychometrics, diet, stool metagenomics, stool consistency, and genome-wide SNP genotypes with autism in 247 Australian children. Surprisingly, the authors suggest their data show a strong correlation between diet and autism spectrum disorder (ASD) but only negligible, if any, ASD-specific microbiome signals. While the first conclusion comes as no surprise, we were rather puzzled by the second conclusion given the growing evidence of strong associations between the microbiome and ASD phenotype and the wide consensus on a close connection between diet and microbiome composition and function. The causal model proposed by Yap *et al*. seemed to imply that diet and the microbiome were two independent variables. A careful review of the approach used by the authors confirmed our suspicions that the statistical models were mis-specified, i.e. they had a questionable biological assumption—the independence of diet and microbiome—embedded in them. We have run side-by-side simulations of the causal linear model proposed by Yap *et al*. and of an analogous model in which diet and the microbiome are treated as co-dependent variables. We show how the Yap *et al*. model can preemptively exclude any potential host-microbe interactions if the diet-microbiome independence assumption is violated. We believe large-scale efforts such as the one described by Yap *et al*. are essential to advance our understanding of the potential role of the microbiome in ASD and other diseases. But these are highly complex systems to analyze and thus ensuring that the statistical methods used are accurate is essential to avoid drawing any potentially misleading conclusions due to subtle causal assumptions propagated by the statistical models themselves.

## Main text

A growing corpus of metagenomic studies revealing strong, albeit inconsistent, variations in microbial diversity in individuals with ASD compared with neurotypicals (Andreo-Martínez et al. 2021; Bezawada et al. 2020; M. Xu et al. 2019), and of clinical studies showing beneficial microbiome interventions in individuals with ASD (Kang et al. 2019), has triggered an interest in addressing the critical gaps in our knowledge of the mechanisms underlying the potential causal relationships between the microbiome and ASD (Saurman et aI. 2020; Svoboda 2020).

The recently published paper by Yap *et al*. is an example of the ever larger and more comprehensive studies that are being performed to try to answer some of the outstanding questions in the field (Yap 2021). By simultaneously sampling and analyzing multiple omics and other biological information levels, scientists are hoping to arrive at a better understanding of the complex interactions among the microbiome, the gastrointestinal system, and the nervous system.

Causal inference on observational data, however, is challenging due to a variety of confounding factors, including experimental design, technical variability, geographical location, and demographic composition, the inherently compositional nature of microbiome datasets, their high dimensionality, their over-dispersion or variability, and the sparse nature of the observed counts. Importantly, most microbiome datasets, including the dataset analyzed by Yap *et al*., originate from cross-sectional studies, single time point studies that do not provide a dynamic picture of change, making any causal inferences related to the etiology of ASD difficult (Imbens and Rubin 2015).

Conversely, however, many statistical models used to analyze such cross-sectional datasets have subtle causal implications built into their structures, thus special care has to be taken to ensure biological and mechanistic accuracy of the models based on prior knowledge of the systems under study. Incorporation of unfounded biological assumptions in a statistical model is referred to in the causal inference literature as model mis-specification. Avoiding such mis-specifications is key to drawing biologically relevant conclusions from complex microbiomes and, by extension, other omics datasets.

The analysis presented by Yap *et al*. exemplifies how a mis-specification ‘hiding in full view’ can dramatically influence the final conclusions of a study. The authors used a type of linear regression model known as linear mixed effects model. The architecture of this type of model inherently assumes that the connectivity and directionality defining the model are reflective of causality. Yap *et al*. used the model architecture previously proposed by Zhang *et al*. (Zhang et al. 2019) (***Fig. 1a****), a model that implicitly assumes diet and microbiome measurements are independent*.

**Figure 1.**
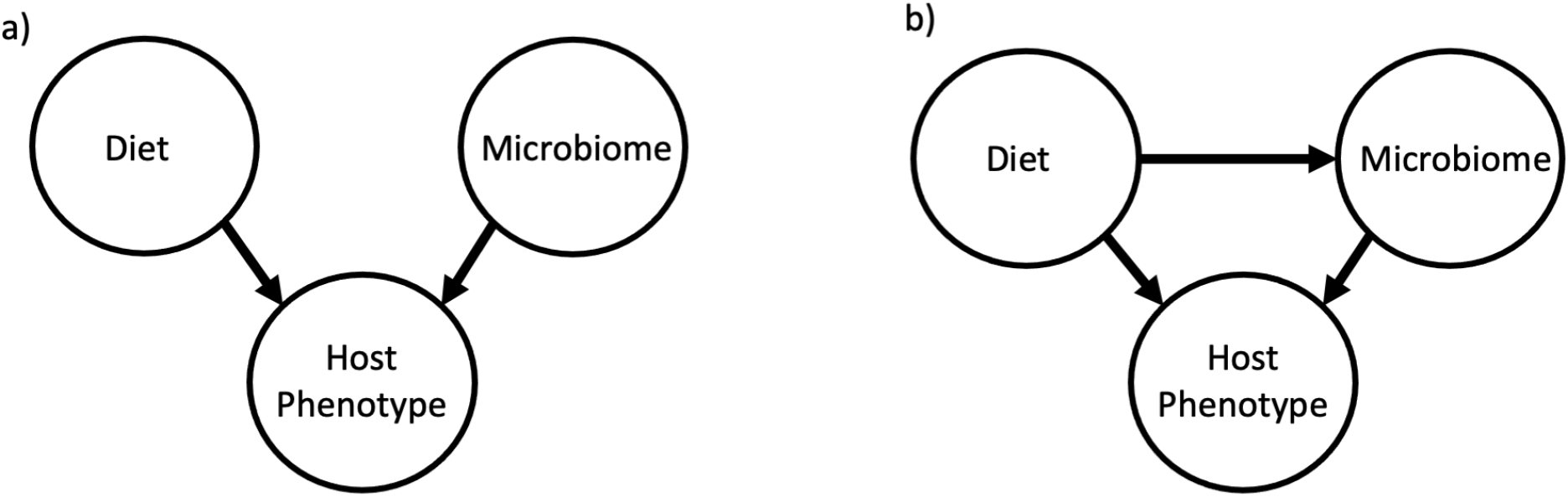
Causal graphs of two linear models illustrating alternate diet, microbiome, and host phenotype interdependencies. *(a) Linear model used in Yap et el*. assuming independent diet and microbiome effects. (b) Modified linear model incorporating the empirically determined co-dependency of diet and microbiome.

This built-in assumption is highly problematic, though, given the lack of evidence for such independence and the overwhelming evidence for the contrary, a co-dependence of diet and microbiome (Singh et al. 2017; Asnicar et al. 2021; Johnson et al. 2020; Turnbaugh et al. 2009). Dietary interventions have been shown to change gut microbiome composition (Wu et al. 2011; David et al. 2014) and microbial by-products have been shown to subsequently enter the bloodstream (Pluznick 2017). A more accurate model would have assumed an architecture that explicitly incorporates the direct influence of diet on the ASD phenotype as well as an indirect influence of diet on the ASD phenotype via the microbiome, *de facto* capturing the known association associations between diet and microbiome composition (**Fig. 1b**).

Model mis-specification can lead to estimation bias (Schisterman et al. 2017). To illustrate the extent to which this bias can affect the outcome of an analysis, we performed a simple side-by-side simulation of the Yap *et al*. model and a modified version of the model that incorporated diet and microbiome data as co-dependent variables (**Fig. 2**; **Suppl. Methods**). In the first scenario, the causal graph shown in Figure 1a is modeled using a linear mixed effects model based on the same causal graph, resulting in a high level of agreement between the estimated and ground truth parameters (**Fig. 2a**). Conversely, if the causal graph underlying the process resembles the causal graph shown in Figure 1b but the data are analyzed using the causal graph shown in Figure 1a to perform inference, then the resulting estimates will show a clear bias (**Fig. 2b**). The more mis-specified the model is, the worse the bias will be. In the above simulation, the ground truth phenotypic variability associated with the microbiome is 83%, however, an analysis using the Yap *et al*. model to estimate the microbiome effect determined that only 3% of the variability could be explained by the microbiome, showcasing how even a strong biological signal can be masked if the model is mis-specified.

**Figure 2:**
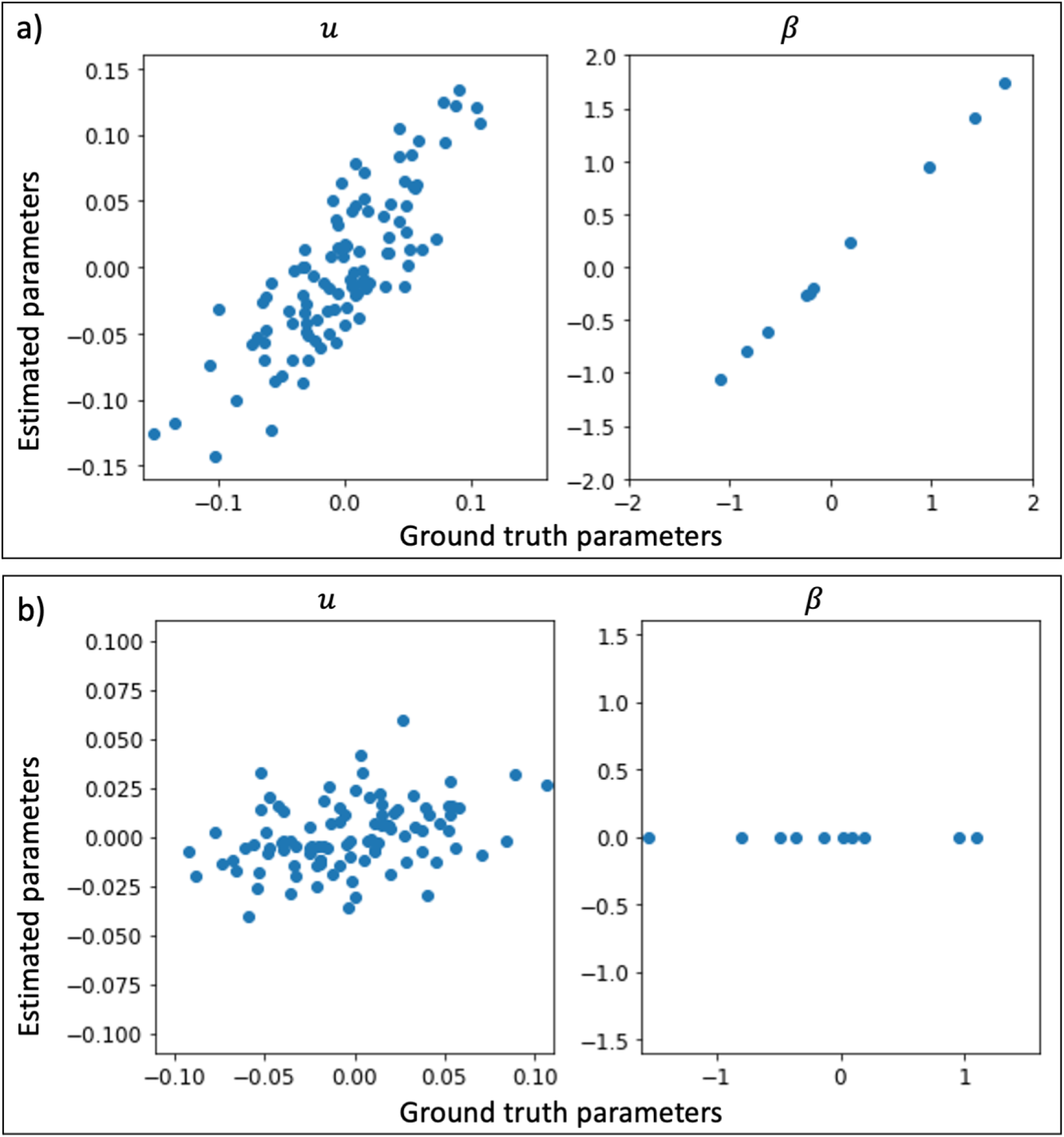
Simulated outcomes of a linear model mis-specification. (a) Comparison of ground truth—*µ* [microbial] and *β* [diet]—and estimated parameters under the independence assumptions of the Yap *et al*. model. (b) Comparison of ground truth and estimated parameters under codependency assumptions.

Causal inference is challenging to perform due to a wide range of confounding factors that may hamper model estimation, and while some studies have suggested that causal inference can be performed on cross-sectional studies using a restrictive set of assumptions (Reichenheim and Coutinho 2010), this remains a hotly contested topic with many in the field subscribing to Donald Rubin’s and Guido Imbens’ assertion that there is “no causation without manipulation” (Imbens and Rubin 2015). Study designs that incorporate longitudinal sampling and interventions make it possible to perform causal inference in the context of Rubin’s and Imbens’ potential outcomes framework. But whether based on cross-sectional or longitudinal and/or interventional datasets, all causal analysis methodologies require model-specific assumptions to be valid in order for the model to be correctly specified. Discussing all the different assumptions is out of the scope of this article, but the topic has been extensively reviewed elsewhere (Imbens and Rubin 2015; Pearl 2000).

While the issue of model assumptions and specifications has not yet been explored in depth in the microbiome field, in other areas, notably in epidemiology, it has long been at the center of discussion due to its profound implications for research and public policy (Rothman and Greenland 2005; Pirracchio, Petersen, and van der Laan 2015; Greenland and Finkle 1995; Vandenbroucke, Broadbent, and Pearce 2016). Given the explosion of reports describing associations and potential causal links between the microbiome and disease, and the emerging role of microbiome data in epidemiological studies, causal inference from microbiome studies needs to take center stage in order to maximize insights and to minimize misinterpretations (Vogtmann and Goedert 2016). We anticipate that appropriate model specifications and careful considerations of the underlying causal assumptions will also help improve reproducibility of microbiome studies.

Based on the evidence we present here, we believe the causal model proposed by Yap *et al*. is acutely mis-specified due to the use of a linear mixed effects model that erroneously assumes independent diet and microbiome effects on the ASD phenotype. Indeed, the main conclusion of the paper, stated in the title of the article as “Autism-related dietary preferences mediate autism-gut microbiome associations”, itself points to the fact that diet does influence the gut microbiome and hence should not be regarded as an independent variable. The dataset analyzed in the paper represents one of the most expansive and comprehensive efforts to date to understand the interplay among different omics levels, including the microbiome, and other parameters in ASD, but we believe the analysis of the data needs to be revisited in order to draw a biologically relevant conclusion with regards to the association of diet and the microbiome with ASD in the 247 individuals who participated in the study. The question of the interplay between diet and the microbiome, and the mechanistic links that might exist to specific ASD genotypes or GI comorbidities is of high priority not only to better understand the etiology of autism, but also to potentially develop therapies designed around particular microbiome manipulations that could help improve the quality of life of both the individuals with ASD and their families. Thoroughly addressing these issues will require deliberate efforts into determining causality that bring together both optimal study design and carefully constructed statistical models that incorporate the most accurate biological assumptions.

## Supplemental Methods

Figure 1a shows the causal directed acyclic graph (DAG) for the following linear model.

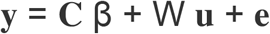

Here, y represents the 0/1 dummy variables representing the ASD phenotype, C represents confounders such as age and sex and diet, β represents the confounder coefficients, W represents clr transformed microbial abundances, u represents a random effects vector and e represents residuals.

As noted in the main text, this model assumes independence between diet and microbes. However, this causal DAG can be modified to encode this relationship given in Figure 1b. This structural equations that encode this model are given by

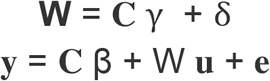

Where *γ* are coefficients that represent the impact of dietary compounds on microbial abundances, and *δ* represents the variability of this impact.

## Notes

### Competing Interest Statement

G.T.-O. is a Consultant-in-Residence at the Simons Foundation

